# Predicting statistics of gene translocation events: role of chromatin compaction and double-strand DNA break

**DOI:** 10.1101/2025.08.11.669613

**Authors:** Anirudh Jairam, Shuvadip Dutta, Sangram Kadam, Kiran Kumari, Ranjith Padinhateeri

## Abstract

How the 3D organization of chromatin affects gene translocation is an open question. By Monte Carlo simulation of translocation events in folded chromatin, we examine how the probability of translocation (*T*_*ij*_) is influenced by chromatin compaction and the occurrence of double-stranded DNA breaks (DSBs). Our results show that, beyond the two-segment contact probability, *T*_*ij*_ depends on the overall compaction of the polymer and has a non-trivial contribution from multi-bead contacts and the probability of DSBs. We propose an empirical formula and provide analytical arguments for simple cases.

---

The genetic code is encoded as a sequence of nucleotides and exist in the form of double-stranded DNA (dsDNA). In living cell, the DNA is further folded with help of a large number of proteins to form a long polymer called chromatin. Every biological processes, including transcription, replication, and stochastic changes in DNA sequence leading to evolution, happen in the context of chromatin [1–3]. The folding of DNA into chromatin not only aids in genome packaging [3, 4] but is also crucial for gene regulation [5, 6] and protecting genes from damage [7–10].

The precise nature of the three-dimensional (3D) organization of chromatin is crucial for many biological processes [11–13]. One experimental approach to determine the 3D organization of the chromatin polymer is chromatin conformation capture [14–16]. A high-throughput version of this method, known as Hi-C, measures the frequency of contact between any two regions of chromatin [17, 18, 21]. These methods have revealed that chromatin exhibits a hierarchical structure, with domains present at various length scales [19, 20]. It has been shown that the probability *C*_*ij*_ of two DNA segments *i* and *j*, along the chromatin contour, coming into contact follows a power-law scaling relation: *C*_*ij*_ ~ |*i −j*| ^*−γ*^ [21, 22]. Depending on the local nature of compaction, *γ* can vary, typically ranging from 2.2 to 0.5 [23–25]. For a randomly folded polymer, *γ* = 3*/*2, while for a chromatin at longer length-scales *γ ≈* 1 [21].

In cells, dsDNA is frequently damaged or broken and undergoes regular repair [20, 26, 27]. Some of the pathways involved in resolving dsDNA damage can lead to genetic recombination or chromosomal translocation [28, 29]. Unlike in meiosis, natural recombination is uncommon in interphase cells [4]. Within the territory of a single chromosome, homologous pairs are rarely in close proximity, making it more likely that breaks induced during interphase are resolved through non-homologous end-joining (NHEJ), which can lead to DNA repair or translocation [30–33]. This suggests that if chromatin segments with multiple DNA breaks are in close 3D proximity, the likelihood of translocation between them increases.

The role of 3D proximity between chromatin segments in facilitating recombination has been investigated using various techniques [34–37], including High-Throughput Genome Translocation Sequencing (HTGTS) [38], which provides translocation frequency information. Lee et al. [39] found a correlation between the spatial proximity of two genomic regions and the likelihood of homology-mediated recombination as a mechanism to repair double-stranded breaks (DSBs). Zhang et al. [36] used *γ*-radiation to induce random DSBs throughout the genome and observed that higher contact frequencies between two regions were associated with increased translocation frequencies in mouse models lacking DNA repair proteins such as ATM. They also found that most translocations in this context were mediated through the Non-Homologous End Joining (NHEJ) repair pathway. Alt et al. [40] proposed that the basic relationship between translocation frequency and contact probability can be expressed as:

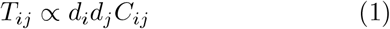

where *T*_*ij*_ and *C*_*ij*_ represent the translocation probability and contact probability, respectively, between genomic regions *i* and *j*, while *d*_*i*_ and *d*_*j*_ denote the DSB frequencies at locations *i* and *j*. However, this equation does not account for the roles of proteins involved in the repair process or for biases that favor self-repair of DSBs over translocations and genomic rearrangements [40]. Additionally, the equation does not incorporate any information about the nature of chromatin organization or the surrounding environment beyond the specific locations *i* and *j*.

Given Eq. 1 a natural way to proceed is *T*_*ij*_ = *kC*_*ij*_*d*_*i*_*d*_*j*_ and assume that *k* is a constant. However, if the proportionality constant depends on the local compaction and other properties, *k* will not be a constant. To investigate this, we generated 3D chromatin configurations and simulated translocation events based on proximity of regions. From the translocation probabilities computed in the simulations, we estimated *k*. Our results show that the overall compaction of chromatin influences the parameter *k* and thereby influences the translocation of two genomic regions.

## Model and Methods

Here, we describe a model to study the relationship between the 3D organization of chromatin and translocation, and a Monte Carlo method to simulate translocation events, in three parts:

(i) Generating an ensemble of single-cell-level inter-chromatin contact information from population-averaged Hi-C data, (ii) Monte Carlo moves to simulate double-stranded DNA breaks and translocation in accordance with 3D chromatin contacts, and (iii) Measurements and analyses based on our simulation data.

We consider chromatin as a linear polymer made of *N* coarse-grained beads [41–50] with each bead representing *l* kb of chromatin. To account for additional contacts due to intra-chromatin interactions, we use population-averaged contact probability matrix *C*_*ij*_ either from Hi-C experiments or from theoretical description of chromatin-like model polymers. We then generate an ensemble of binary contact matrices (*M*_*ij*_), each representing a single chromatin polymer configuration. To achieve this, we perform the following procedure. Choose a pair of beads (*i, j*) from a single polymer and assign a contact between the pairs if *r*_*n*_ *< C*_*ij*_, where *r*_*n*_ is a random number drawn from a uniform distribution of numbers in the range [0,1]. If the contact is assigned, the corresponding element in a matrix is set to *M*_*ij*_ = 1, and *M*_*ij*_ = 0 otherwise. Repeating this for every possible pair once gives us contact information of a single configuration of chromatin.

Given the *M*_*ij*_ matrices representing chromatin configurations, we implement Monte Carlo moves resulting in DSBs and translocation. A DSB at location *i* is considered as breaking of a DNA segment, belonging to the *i*^*th*^ coarse-grained bead, with a probability *d*_*i*_. This results in a 1D binary vector (*b*_*i*_ = 0 or 1) representing whether the bead is broken in a given chromatin polymer.

The broken beads are fixed as follows: we choose a broken bead (*i*) randomly (i.e., *b*_*i*_ = 1), and find out all other broken beads which are in contact with the chosen bead. That is, for the chosen bead *i*, find all matrix elements where *M*_*ij*_ = 1 as well as *b*_*j*_ = 1. If there are *N*_*b*_ broken beads in contact with *i* (including *i*), we “repair” the breakage with one of them randomly (i.e, with probability 1*/N*_*b*_). If the broken bead *i* chooses to repair with any other bead *j*, we call it a translocation between *i* and *j*. If a segment chooses itself (*i* choosing *i*), then this would be called a DSB repair and we don’t count this outcome as a translocation. We repeat this process until all broken segments are either self-repaired or translocated, completing one simulation realization. When two DSBs occur, there are four broken ends. We assume that when one pair of broken ends are translocated, the remaining pair will translocate with each other. The whole process of breakage and repair is repeated *™*10^8^ *−* 10^9^ times ensuring that we cover the entire space of outcomes and sample enough translocations to generate good statistics to compute the probability of translocation between segments *i* and *j, T*_*ij*_ defined as

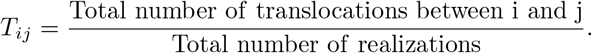

Prompted by the experimental observations by Chiarle et al. [38], we assume that all translocations in our simulations follow NHEJ. Hence, there is no involvement of DNA sequence as in the case of Homology-directed repair. In this work, we assume that all the repair/translocation events are stochastic and any effect due to proteins will only affect the translocation timescale, which is not relevant in this study as we are investigating only the steady-state properties. All possible scenarios of rearrangements, including deletions and circularization of small segments, are accounted in our procedure. Once we obtain *T*_*ij*_, we define the translocation fraction as:

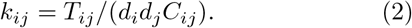

Here, *k*_*ij*_ represents the ratio of the actual probability of translocation to the expected translocation probability from an existing simple theory (see Eq. 1). We also compute the “mean translocation fraction” defined as: 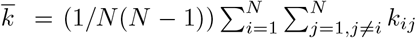. Throughout this work, we assume *d*_*i*_ = *d*_*j*_ = *d*. In this study, we present translocation statistics computed using various polymer models, including: (i) a Random Walk (RW) polymer having *C*_*ij*_ *∝* | *i −j*| ^*−*3*/*2^, (ii) a Self Avoiding Walk(SAW) polymer, (iii) polymer globules where the chromatin is considered as a bead-spring chain with every bead having a uniform attractive interaction with every other bead via a Lennard-Jones (LJ) potential, (iv) a region of IMR90 and K562 cell lines based on coarse-grained contact maps as simulated by Kumari et al. [43]. We also used the contact maps of *Candida tropicalis* genome from published data [51].

## Results

We present results from our translocation simulations and examine how DSB and contact probabilities influence the translocation probability. The null expectation is that *k*_*ij*_ = *k*, a constant independent of (*i, j*). We will show that this expectation does not hold and *k*_*ij*_ is influenced by multiple parameters, given the statistical picture of translocation.

### k_ij_ is not a constant but a function of chromatin compaction

To understand the role of chromatin compaction on *k*_*ij*_, we simulated the following polymers (*N* = 50) with different contact probability matrices: these include Random Walk (RW), Self Avoiding Walk(SAW), different types of polymer globules(see SI), and regions from IMR90 and K562 cell lines [43].

Starting with a contact matrix *C*_*ij*_ for each polymer, we introduced DSB and performed DSB repair/translocation as described in Methods. We calculated *k*_*ij*_ using Eq. 2. Fig. 2a shows that the *k*_*ij*_ is not a constant but has a distribution. We find that both the mean and the nature of the distribution depend on the type of the polymer. Highly packed polymers such as the compact globule (black) show lower average *k*_*ij*_ compared to open structures like the SAW (blue) or RW(pink). We also observe a slight difference in the *k*_*ij*_ distributions of SAW and RW polymer consistent with the fact that RW is slightly more compact than SAW. We have simulated globules in a few different ways and all of them have *k*_*ij*_ values smaller than SAW and RW. Chromatin structures from the two cell lines (IMR90 and K562) yield *k*_*ij*_ values that lie between these extremes. Notably, it is known that the chromatin region studied is slightly more compact in IMR90 than in K562 [43] and we observe a corresponding difference in the mean *k*_*ij*_ values. All of these results indicate that chromatin compaction influences *k*_*ij*_ and hence translocation probabilities. We define mean contact probability 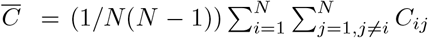 as a measure of compaction and examine its relation with *k*_*ij*_. Since we find an inverse dependence, we plot 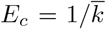 versus 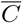 showing a linear dependence (see Fig. 2b). This quantifies how 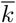 varies for different polymers in Fig. 2a. The intercept in Fig. 2b is 2; this is because, for a translocation to occur, there has to at least two choices — repairing the DSB or translocation among the two broken pairs. By fitting straight lines in Fig. 2b, we obtain the following empirical relation:

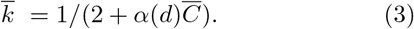

where *α*(*d*) is the slope which is a function of DSB probability *d*.

**FIG. 1.**
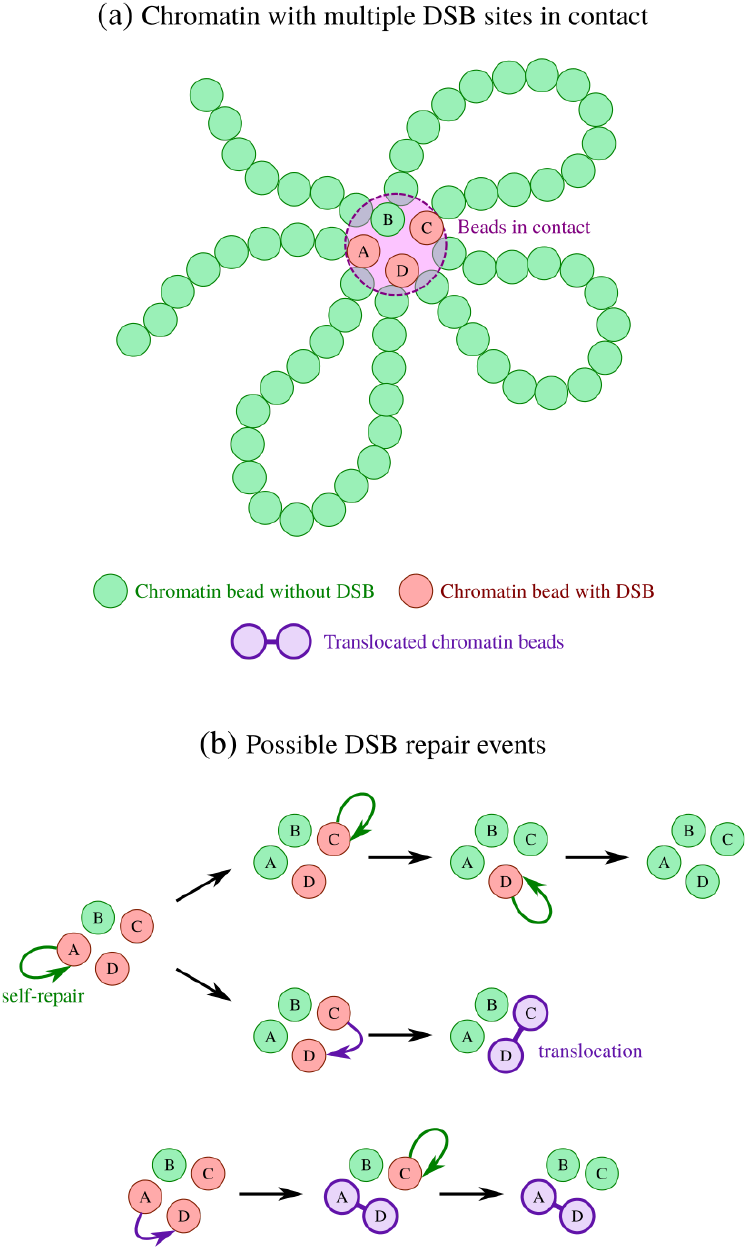
(a) A polymer configuration with 4 segments (beads) in contact (see circle); 3 among them contain DSBs. (b) We simulate various possible DSB repair and translocation events. Top trajectory: segment-A, segment-C and segment-B get repaired with themselves one after the other. Middle trajectory: after the self-repair of segment-A, segments B and C get translocated with each other. Bottom trajectory: segment-A and D get translocated. Segment-C repairs with itself.

**FIG. 2.**
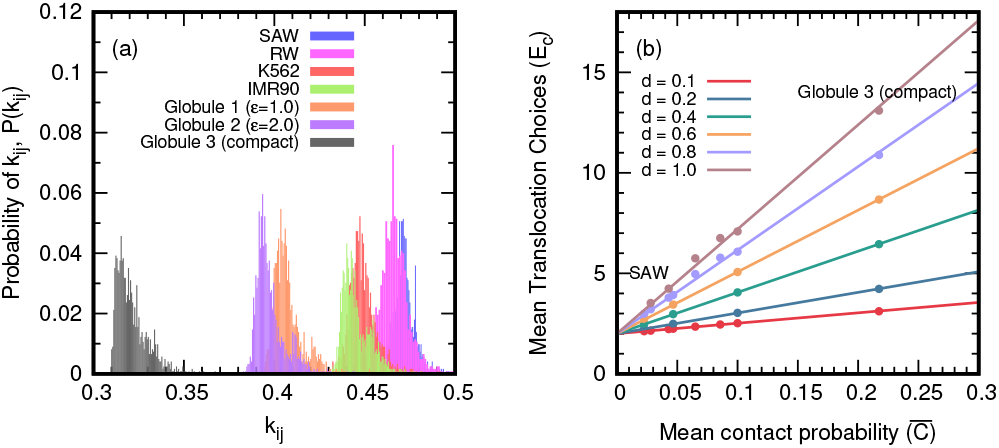
Contrary to the prevalent notion, the translocation fraction (*k*_*ij*_) is a function of polymer compaction: (a) Distribution of *k*_*ij*_ for different polymers in the inscreasing order of compaction: SAW, RW, chromatin regions from K562 and IMR90 cell lines, 3 different globule-like polymers with different conpaction (see SI). Here, *d* = 0.1. (b) 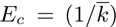 linearly increases with the mean contact probability 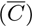 for the same polymer chains in (a).

### The Translocation fraction varies with DNA double-strand break probabilities

To understand the dependence of translocation fraction (*k*_*ij*_) on DSBs, we varied DSB probability (*d*) for a given contact probability matrix. Fig. 3 shows how *k*_*ij*_ varies for two very different polymers, SAW (Fig. 3a) and the compact globule (Fig. 3b) for various values of *d*. Assuming that each segment breaks with an equal probability *d* we show that as *d* increases, *k*_*ij*_ decreases. Since increasing *d* increases the number of possible choices for a segment *i* to translocate, the fraction of translocation with any particular segment *j* will fall. Hence the peak values of *k*_*ij*_ and mean values 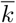 will depend on *d* and compaction.

**FIG. 3.**
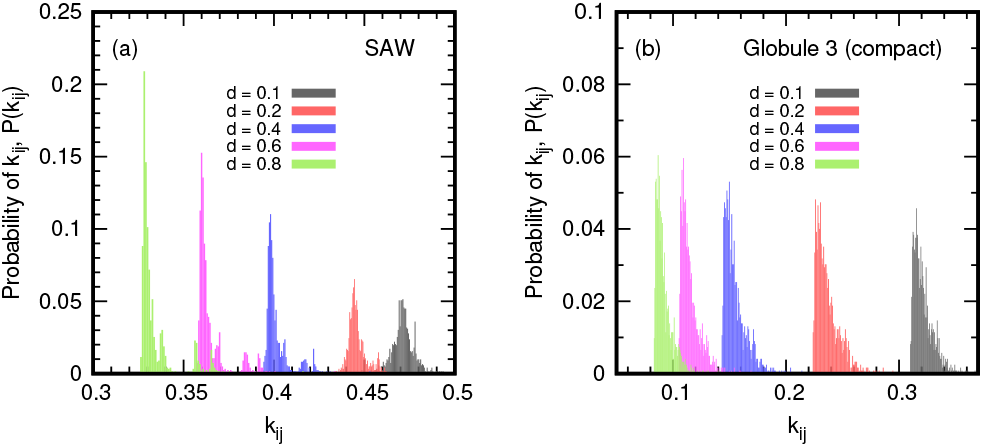
Translocation fraction (*k*_*ij*_) is a function of DSB probability (*d*): Distribution of *k*_*ij*_ values from (a) SAW and (b) compact globule for various *d* values. Peak values of *k*_*ij*_ reduce much less for higher *d* in the SAW as compared to the compact globule due to its relatively more open structure.

The observation that 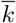 is a function of the double strand break probability *d* implies that *T*_*ij*_ is no longer proportional to *d*^2^ but some other function of *d*. In Fig. 4 we plot the mean of 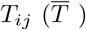 as a function of *d* showing the deviation from the *d*^2^ behavior. For a polymer like the compact globule, there is a larger deviation of 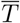 from the *d*^2^ behavior as compared to a RW polymer. As compact polymers have many contacts per segment, the limiting parameter for translocation is not the number of available contacts, but the available DSBs determined by the parameter *d*. This implies that increasing *d* will rapidly reduce 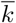 for such polymers (see Fig. 3). In the case of open polymers, the limiting parameter is not *d* but the average number of contacts. Increasing *d* does little to increase the mean translocation choices (*E*_*c*_) of the system. Hence, the dependence of their 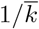 values on *d* is weaker. We also see that *E*_*c*_ increases linearly with increasing *d* (Fig. S1) as 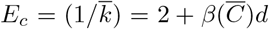, suggesting,

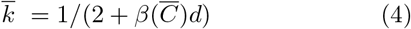

where 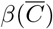is the slope for a particular polymer type.

**FIG. 4.**
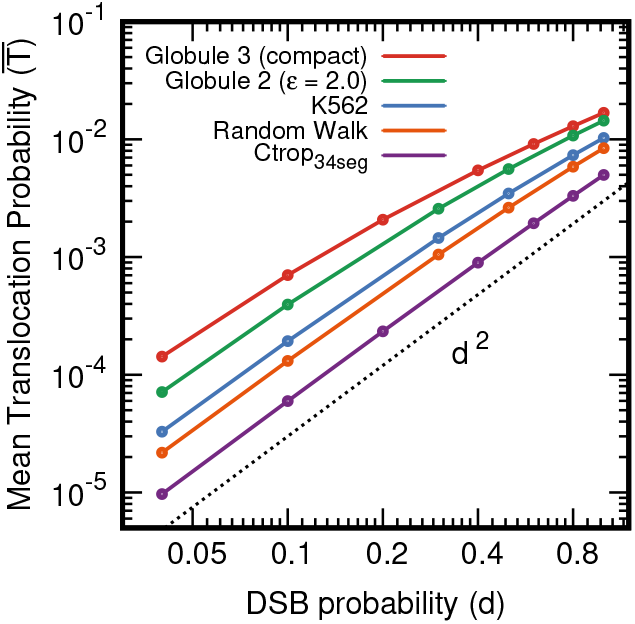
*T*_*ij*_ need not be proportional to *d*^2^ unlike the prevalent notion. Since 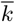 is a function of *d*, the translocation probability cannot be a function of *d*^2^. The relatively open polymer RW is closer to the *d*^2^ powerlaw while more compact polymers deviate from *d*^2^.

### Analytical expressions for T_ij_ and k_ij_

When we consider a polymer with two segments *i* and *j*, the only translocation possible is between *i* and *j*, and the probability is *T*_*ij*_ = *k*_*ij*_*C*_*ij*_*d*_*i*_*d*_*j*_. As mentioned earlier, for the two segment system 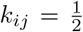. However, when one adds a third segment *m*, it is possible to have a breakage in segment *m* with probability *d*_*m*_ and contacts leading to translocations between (*i, m*) and (*j, m*). Accounting for all these possibilities (see SI) we find that the translocation probability for a three segment system is given by

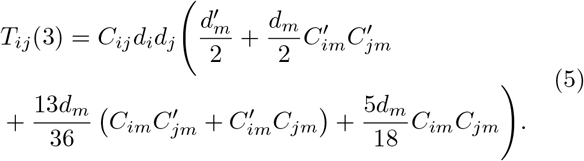

Here *C*_*im*_ is the contact probability between segments *i* and *m*, and *C*_*jm*_ is the contact probability between segments *j* and *m*. 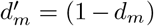 is the probability that segment *m* is not broken. The 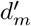 term represents the cases where only *i* and *j* are broken. The second term represents the case where *m* is broken but neither *i* nor *j* is in contact with *m* with 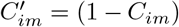 and 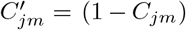. The third term represents the case where the segment *m* is broken but only one of the pairs involving *m* (either *im* or *jm*) is in contact. The last term represents the case where all 3 segments are broken and in contact with each other. The counting of all possible combination of these gives the numerical pre-factors as explained in SI. Simplifying by substituting for the primed quantities gives us:

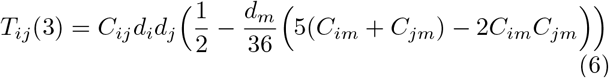

This is equivalent of defining

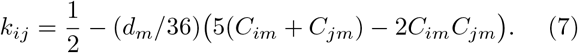

We also find that the maximum value of *k*_*ij*_ (upper limit) can be 1*/*2, which occurs when *d*_*m*_ *→* 0 or *C*_*im*_ *→* 0 and *C*_*jm*_ *→* 0. We performed a translocation simulation for a three segment system and obtained mean values of *k*_*ij*_ to show that our analytical relation (Eq. 7) matches well with the simulation results (Fig. 5a) for simple cases where *C*_*im*_ = *C*_*jm*_=constant.

**FIG. 5.**
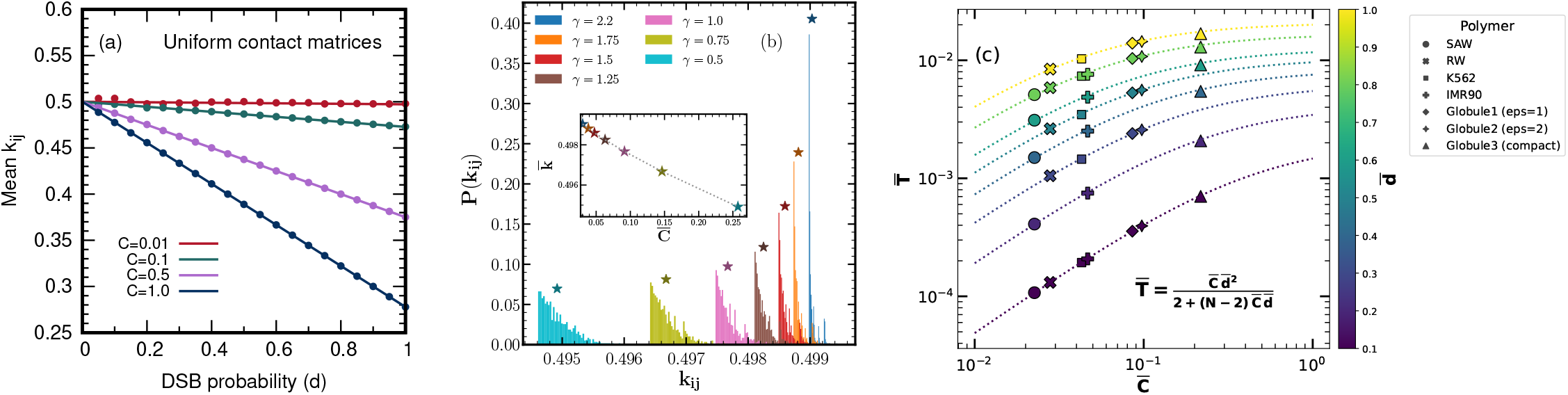
(a) Mean 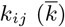 computed from the three-segment model as a function of DSB probability for a 3 *×* 3 contact matrix having all non-diagonal elements as 0.01, 0.1, 0.5 or 1.0. The solid lines are plotted using Eq. 7 and the points are generated from simulations. (b) Distribution of *k*_*ij*_ = *T*_*ij*_*/*(*C*_*ij*_*d*_*i*_*d*_*j*_) from Eq. 8, with *C*_*ij*_ = |*i −j* |^*−γ*^ and *d* = 3*/N*, for various *γ* values. Asterisks mark the mean value 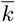. Inset: 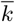 vs. 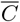 from the analytical theory reveals that higher chromatin compaction (larger 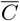) leads to reduced 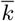, highlighting the role of multi-body contacts in *T*_*ij*_ estimates. (c) Average translocation probability 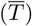 as a function of mean contact probability 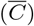 from the Eqs 10 and 11 with 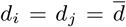 (see colorbar) for *N* = 50. Points represent the 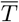 calculated from the simulation for different polymers as described in Fig. 2a.

We can extend the analytical relation in Eq. 6 for the case of a polymer with *N* segments by expanding the *T*_*ij*_ in terms of the number of breaks. In principle, the expansion will have terms up to *𝒪* (*d*^*N*^). However, if we assume that four or higher breaks are improbable as well as higher order multi-bead contacts are improbable, we can re-write the equation as follows:

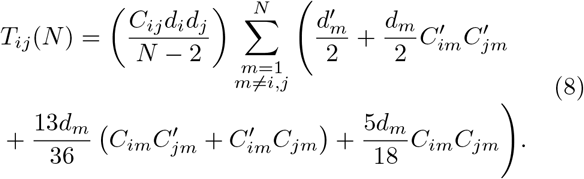

Here two out of these three breaks should be on *i* and *j* so that we can compute *T*_*ij*_; however, the third break can be on any of the other *N −* 2 beads. We computed the *T*_*ij*_(*N*) from the above equation by taking *C*_*ij*_ =| *i −j* |^*−γ*^ and *d* = (3*/N*). Using Eq. 2, we get *k*_*ij*_ and its distribution is plotted in Fig. 5b for several *γ* values. As the compaction increases (as *γ* decreases), the peak of the distribution shifts away from 0.5, as predicted by our simulations. Here the trend is similar to what is observed in Fig. 2a, where full simulations without approximation produce *k*_*ij*_ distributions that shift away from 0.5 as the compaction increases. We emphasise that the goal of this subsection is not to determine the exact functional form of *k*_*ij*_, but rather to show analytically that *k*_*ij*_ is not constant and indeed varies with compaction.

We can also derive a lower bound for *k*_*ij*_ in a theoretical limit when all segments are broken (*d*_*i*_ = 1 for all *i*) and and all segments are in contact with each other (*C*_*ij*_ = 1 for all *i* and *j*). For an *N*-segment polymer, we derive a recursive relation for the lower bound of *k* as

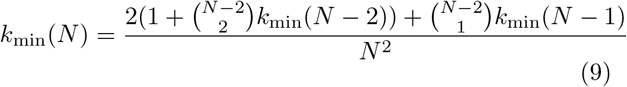

where *k*_min_(*N −*2) and *k*_min_(*N −*1) are the lower limits of 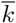 for an *N −* 2 and *N −* 1 long polymers respectively (see SI). *k*_min_(2) and *k*_min_(3) are 0.5 (2 choices on average) and 0.2778 (minimum value of *k*_*ij*_), respectively. It turns out that 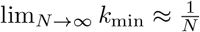 (Fig. S2). Using the relations 3, 4 and 9 we can obtain an empirical formula for the mean value 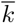 as:

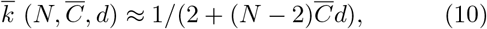

and the corresponding translocation probability as

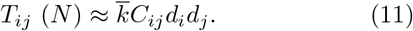

We show that this empirical formula matches well with the data we obtained from our simulations for different polymers (Fig. 5c).

One important insight from this study is that regions with more contacts, like heterochromatin regions or TAD boundaries, will have low *k*_*ij*_ values. This can be seen in Fig. S3 and shows that TAD boundaries have lesser *k*_*ij*_ values as compared to other regions. An interesting notion emerging from the results of these simulations is that, due to excessive packaging in compact regions, each segment has many ‘decoy’-like segments to which it can potentially translocate. This reduces the probability of important sites (active promoters, enhancers) translocating or moving to undesirable segments. In this way, harmful translocations may be suppressed by chromatin packaging, in addition to other biological mechanisms. Whether this effect is biologically significant remains unknown.

One drawback of this model is that it does not provide information about what happens after translocations occur. That is, we likely cannot use the same contact matrix after one realization if we wish to study the time evolution of translocation probabilities. Also, the model takes into account all translocation/rearrangement events including deletions and circularization which may not be favorable for species and may not be seen in living populations. Eq. 11 only stands when all events are purely statistical.

To conclude, we investigated how the 3D polymer organisation affects the translocation probability. We simulated different types of polymers with different 3D organization and compaction. From the simulation, we obtained a functional relation between translocation probability, contact probability and double-strand DNA break probability. Naive expectation from the simplest possible model is that *T*_*ij*_ *∝ d*_*i*_*d*_*j*_*C*_*ij*_, as shown in Eq. 1. However, we show that translocation probability depends not only on the pairwise contact probability *C*_*ij*_ but also exhibits a non-trivial dependence that reflects the multi-bead contacts, the overall compaction of the polymer, and the DSB probability. Using our simulation data, we derived an empirical relation for the mean translocation probability that is particularly relevant under conditions of high DSB rates and high chromatin compaction.

## Acknowledgement

RP acknowledges funding from Science and Engineering Research Board (SERB) (Grant number: CRG/2022/007974) and Sunita Sanghi Centre of Aging and Neurodegenerative Diseases (SCAN), IIT Bombay. SD acknowledges fellowship support from the PrimeMinister’s Research Fellowship, Ministry of Education, India.

