## Supplementary Information for "Predicting statistics of gene translocation events: role of chromatin compaction and double-strand DNA break"

(Dated: August 10, 2025)

### S1. POLYMER MODELS

We have used a bead-spring polymer model consisting of N beads each of size (diameter) $\sigma$. The total energy of the polymer is given by

$$E = \sum_{i=1}^{N-1} E_{spring}(|\vec{r}_{i+1} - \vec{r}_i|) + \sum_{i=1}^{N-1} \sum_{j=i+1}^{N} E_{LJ}(|\vec{r}_j - \vec{r}_i|) \tag{S1}$$

Here $\vec{r}_i$ is the position vector of $i^{th}$ bead. The first term represents the polymer connectivity through harmonic springs given as

$$E_{spring}(r) = \frac{k}{2}(r - r_0)^2 \tag{S2}$$

Here $r_0$ is the equilibrium distance of the spring, $r$ is the 3D distace between polymer beads and $k$ is the spring constant. The value of spring constant is $k = 10$ and $r_0 = 1.0$. The second term in Eqn. S1 represents the attractive interactions through Lennard-Jones potential given by,

$$E_{LJ}(r) = \begin{cases} 4\epsilon\left[\left(\frac{\sigma}{r}\right)^{12} - \left(\frac{\sigma}{r}\right)^{6}\right] & r < r_c, \\ 0 & r \geqslant r_c. \end{cases} \tag{S3}$$

Here $\epsilon$ is the energy parameter that determines the strength of the attractive interaction, $\sigma$ is the diameter of the bead and $r_c$ is the cutoff for the potential.

#### Simulation scheme

The Langevin dynamics simulations were performed using the LAMMPS software package [1]. In this approach, the position of each polymer bead evolves according to the equation:

$$m_i \frac{d^2\mathbf{r_i}}{dt^2} = -\nabla E_i(\mathbf{r}) - \gamma_i \frac{d\mathbf{r_i}}{dt} + \sqrt{2\gamma_i k_B T}\boldsymbol{\eta_i}(t) \tag{S4}$$

where $r_i$ is the position of the center of mass of the $i$-th bead, $m_i$ is its mass, $E_i$ is the sum of all the interaction potentials acting on bead $i$. The friction coefficient $\gamma_i$ determines the diffusion constant $D_i = k_B T/\gamma_i$. For polymer beads, we set $m_i = 1$ and $\gamma_i = 1.0$ for polymer beads. $\eta_i(t)$ is a noise term with components which satisfy: $\langle \eta_{i\alpha}(t) \rangle = 0$ and $\langle \eta_{i\alpha}(t)\eta_{j\beta}(t') \rangle = \delta_{ij}\delta_{\alpha\beta}\delta(t - t')$, where $\eta_{i\alpha}(t)$ is component $\alpha$ of the noise vector for object $i$, and $\delta_{ij}$ and $\delta(t)$ are the Kronecker and Dirac delta functions respectively. These equations are solved using a velocity-Verlet algorithm with time step $dt = 0.005\tau$, where $\tau$ is the simulation time unit defined by $\tau = (m\sigma^2/k_B T)^{\frac{1}{2}}$.

*Electronic address:
†Electronic address:
‡Electronic address:

**Parameters for Different Polymer Simulations**

1. **Self-Avoiding Walk (SAW):** In the SAW model, the Lennard-Jones (LJ) interaction strength was set to $\epsilon = 1.0$, with a cutoff radius of $r_c = 2^{1/6}\sigma$ to ensure purely repulsive interactions.
2. **Random Walk (RW):** In the RW model, the interaction strength was set to $\epsilon = 0.0$, effectively eliminating LJ interactions and necessarily simulating a Gaussian chain.
3. **Globule-1 and Globule-2:** For the two globular configurations, the LJ interaction strengths were set to $\epsilon = 1.0$ (Globule-1) and $\epsilon = 2.0$ (Globule-2), respectively. A cutoff radius of $r_c = 2.5\sigma$ was used in both cases to include attractive interactions.
4. **Globule-3 (Compact Globule):** For the compact globule (Globule-3), the LJ interaction strength was set to $\epsilon = 2.0$ with a cutoff radius of $r_c = 2.5\sigma$. To increase the average contact probability per segment, LJ interactions between adjacent segments ($i$ and $i+1$) were excluded.

### S2. DERIVATION OF $k_{ij}$ FOR A THREE-SEGMENT SYSTEM

Consider a polymer system composed of three segments, labeled $i$, $j$, and $k$. This system is characterized by six parameters: the contact probabilities $C_{ij}$, $C_{jk}$, $C_{ik}$ between each pair of segments, and the probabilities $d_i$, $d_j$, $d_k$ that each segment is broken. Suppose we are interested in computing the translocation probability $T_{ij}$, which denotes the probability that segments $i$ and $j$ translocate. For translocation to occur between these two segments, both must be broken and must be in contact. We consider all scenarios in which either segment $i$ selects $j$, or $j$ selects $i$, for translocation. The third segment, $k$, may or may not be broken or in contact with the others. We discuss the possible cases in detail below.

#### Case I: Segment $k$ is not broken

In this scenario, segment $k$ is not broken. We consider all configurations of pairwise contacts involving $k$, including whether contacts $C_{ik}$ and $C_{jk}$ are present. Since $k$ is not broken, it cannot participate in translocation events. Thus, the probability that $i$ and $j$ translocate together is:

$$k_{ij}(\mathrm{DSB}=2) = \frac{d'_k}{2} \tag{S5}$$

where $d'_k = 1 - d_k$ is the probability that $k$ is not broken. The factor of $1/2$ arises as follows:

1. There is a $1/2$ probability that $i$ is selected first, and it chooses $j$.
2. Similarly, there is a $1/2$ probability that $j$ is selected first, and it chooses $i$.

Averaging over these two possibilities yields a net probability of $\frac{1}{2}$.

#### Case II: Segment $k$ is also broken

Here, all three segments ($i$, $j$, and $k$) are broken, and the probability $d_k$ appears as a common multiplicative factor in all contributions.

##### *Case IIa: Only One Additional Contact Exists*

Assume that only one of the two contacts involving $k$ is present; that is, either $C_{ik} = 1$ and $C_{jk} = 0$, or vice versa. Then, the probability that $i$ and $j$ undergo translocation is:

$$k_{ij}(\mathrm{DSB}=3,\ \mathrm{Contact}=2) = \frac{13}{36}\left[C_{ik}(1-C_{jk}) + (1-C_{ik})C_{jk}\right] \tag{S6}$$

The prefactor $\frac{13}{36}$ is derived as follows:

1. There is a 1/3 chance that any of the three segments ($i$, $j$, or $k$) is selected first.
2. If $i$ is selected first, it chooses among the three segments uniformly; the probability of choosing $j$ is 1/3.
3. If $j$ is selected first, and it is only in contact with $i$, it chooses $i$ with probability 1/2.
4. If $k$ is selected first and is not in contact with both $i$ and $j$, it picks itself. Subsequently, $i$ and $j$ must be selected and choose one another, which occurs with probability $\frac{1}{3} \times \frac{1}{2}$.

Summing these contributions:

$$\frac{1}{3}\left(\frac{1}{3}+\frac{1}{2}+\frac{1}{2}\cdot\frac{1}{2}\right)=\frac{13}{36}$$

*Case IIb: All segments are mutually in contact*

If all three contacts are present ($C_{ij} = C_{ik} = C_{jk} = 1$), then the probability of translocation between $i$ and $j$ is:

$$k_{ij}(\text{DSB} = 3,\ \text{Contact} = 3) = \frac{5}{18} C_{ik} C_{jk} \tag{S7}$$

The factor $\frac{5}{18}$ is obtained by considering the following:

1. Each segment has a 1/3 chance of being selected first.
2. If any segment is selected first, it chooses among the other two with probability 1/3.
3. If $k$ is selected first, it may pick itself, and $i$ and $j$ must then translocate. The combined probability of such events is:

$$\frac{1}{3}\left(\frac{1}{3}+\frac{1}{3}+\frac{1}{3}\cdot\frac{1}{2}\right)=\frac{5}{18}$$

**Final Expression for $T_{ij}$**

Combining the contributions from the above cases, and introducing the notation $C'_{ik} = 1 - C_{ik}$, $C'_{jk} = 1 - C_{jk}$, and $d'_k = 1 - d_k$, the full expression for the translocation probability between segments $i$ and $j$ is:

$$T_{ij} = C_{ij} d_i d_j \left(\frac{1}{2} - \frac{d_k}{36}\left[5(C_{ik} + C_{jk}) - 2C_{ik}C_{jk}\right]\right) \tag{S8}$$

This framework can similarly be applied to compute the translocation probabilities $T_{ik}$ and $T_{jk}$. Taking the average over all three pairwise translocation probabilities yields:

$$\overline{T} = \frac{1}{3}\left(T_{ij} + T_{ik} + T_{jk}\right) \tag{S9}$$

### S3. DERIVING $k_{\min}$ FOR A POLYMER OF SIZE $N$

Here, we derive an expression for the minimum value of the translocation fraction, $k_{\min}$, for a polymer consisting of $N$ segments. Since $(1/k)$ is linearly proportional to both $C$ and $d$, the minimum value of $k$ corresponds to the maximum values of $C$ and $d$, i.e., $C = d = 1$. This scenario represents the case where all segments are broken and are in mutual contact. We now calculate the probability that any two segments $i$ and $j$ choose each other to translocate.

**Case I: Direct selection of $i$ and $j$**

If either segment $i$ or $j$ is selected first (with probability $1/N$), and it selects the correct partner (i.e., $j$ or $i$, respectively, also with probability $1/N$), the total probability of this event is:

$$\frac{1}{N} \cdot \frac{1}{N} + \frac{1}{N} \cdot \frac{1}{N} = \frac{2}{N^2}.$$

**Case II: Another segment $l \neq i, j$ is selected first**

If a different segment $l$ (where $l \neq i, j$) is selected first, two outcomes are possible depending on whether $l$ fixes itself or translocates with another segment that is not $i$ or $j$.

**Case IIa: $l$ fixes itself**

If $l$ selects itself (with probability $1/N$), and there are $\binom{N-2}{1}$ such choices, the probability of this event is:

$$\frac{\binom{N-2}{1}}{N^2}.$$

In this case, $l$ is removed from the pool of broken segments, reducing the system to $N-1$ segments. The minimum translocation probability is then $k_{\text{min}}(N-1)$, and the contribution to $k_{\text{min}}(N)$ becomes:

$$\frac{\binom{N-2}{1} \cdot k_{\text{min}}(N-1)}{N^2}.$$

**Case IIb: $l$ translocates with another segment (excluding $i$ and $j$)**

If $l$ pairs with another segment other than $i$ or $j$, the number of such pairs is $\binom{N-2}{2}$. Since two segments are removed from the broken set, the translocation probability becomes $k_{\text{min}}(N-2)$, and the contribution is:

$$\frac{2 \cdot \binom{N-2}{2} \cdot k_{\text{min}}(N-2)}{N^2}.$$

Combining all contributions, we arrive at the recursive expression for $k_{\text{min}}(N)$:

$$k_{\text{min}}(N) = \frac{2}{N^2} + \frac{\binom{N-2}{1} \cdot k_{\text{min}}(N-1)}{N^2} + \frac{2 \cdot \binom{N-2}{2} \cdot k_{\text{min}}(N-2)}{N^2}. \tag{S10}$$

As translocation requires atleast a pair of broken beads, $k_{\text{min}}(0) = 0$ and $k_{\text{min}}(1) = 0$. Then the recursion yields values such as $k_{\text{min}}(2) = 0.5$ and $k_{\text{min}}(3) = 0.2778$, which are consistent with the simulation results.

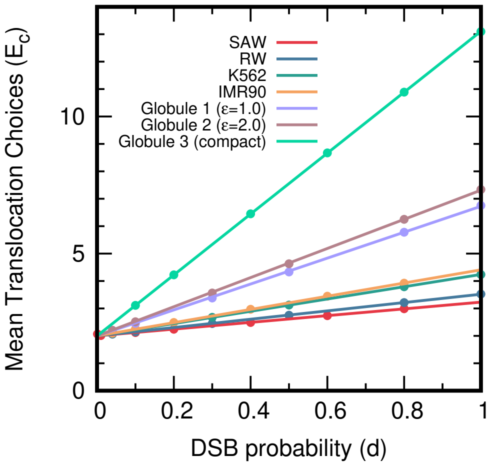


FIG. S1: Mean Translocation Choices ($E_c$) as a function of double-strand break probability ($d$). The value of $E_c(= 1/\bar{k})$ increases linearly with $d$, with the rate of increase (slope $\beta(C)$) depending on the degree of polymer compaction. Among the polymer conformations considered, the Compact Globule exhibits the highest slope, whereas the self-avoiding walk (SAW) conformation shows the lowest slope.

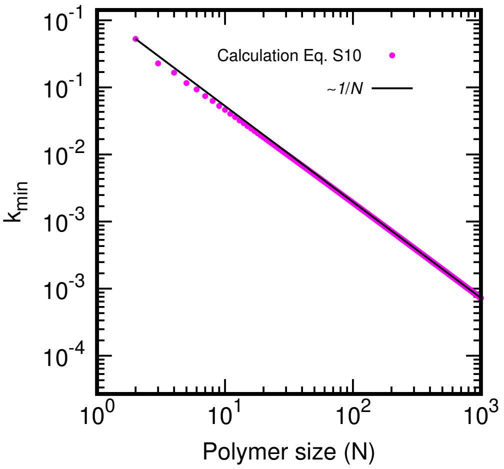


FIG. S2: The minimum translocation fraction, denoted as ($k_{\min} = k_{ij}|_{C=d=1}$). $k_{\min}(N)$ is calculated using $k_{\min}(N) = \frac{2}{N^2} + \frac{\binom{N-2}{1}\cdot k_{\min}(N-1)}{N^2} + \frac{2\cdot\binom{N-2}{2}\cdot k_{\min}(N-2)}{N^2}$. In the large $N$ limit, $k_{\min}(N)$ approximates to $\frac{1}{N}$.

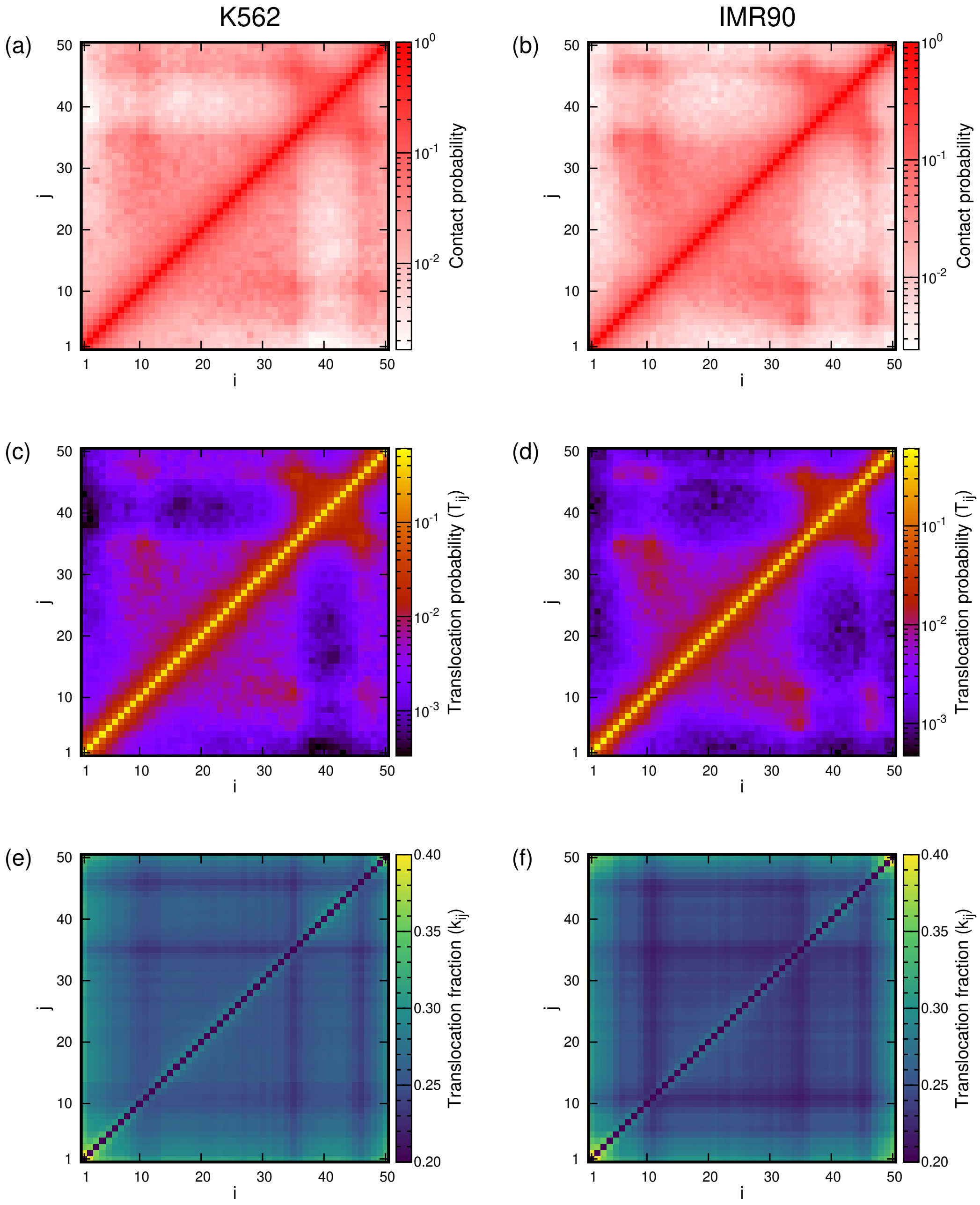


FIG. S3: Comparison of contact probability ($T_{ij}$) and translocation fraction ($k_{ij}$) heatmaps. (a,b) Contact probability matrices for a representative TAD region in the K562 and IMR90 cell lines, respectively. (c,d) Corresponding translocation probability matrices ($T_{ij}$). (e,f) Corresponding translocation fraction matrices ($k_{ij}$). TAD boundaries display lower $k_{ij}$ values compared to intra-TAD regions, supporting the idea that polymer compactionand consequently the number of contactsplays a key role in determining translocation probability..

[1] A. P. Thompson, H. M. Aktulga, R. Berger, D. S. Bolintineanu, W. M. Brown, P. S. Crozier, P. J. in 't Veld, A. Kohlmeyer, S. G. Moore, T. D. Nguyen, et al., Comp. Phys. Comm. **271**, 108171 (2022).
